# Ultrastructural viscoelastic behavior of fibrillar collagen identified by AFM Nano-Rheometry and direct indentation

**DOI:** 10.1101/2024.10.19.619231

**Authors:** Meisam Asgari, Elahe Mirzarazi, Ryan J. Benavides, Yuri M. Efremov, Robert D. Frisina, Hojatollah Vali, Horacio D. Espinosa

## Abstract

Soft tissues exhibit predominantly time-dependent mechanical behavior critical for their biological function in organs like the lungs and aorta, as they can deform and stretch at varying rates depending on their function. Collagen type I serves as the primary structural component in these tissues. The viscoelastic characteristics of such tissues, stemming from diverse energy dissipation mechanisms across various length scales, remains poorly characterized at the nanoscale. Prior experimental investigations have predominantly centered on analyzing tissue responses largely attributed to interactions between cells and fibers. Despite many studies on tissue viscoelasticity from scaffolds to single collagen fibrils, the time-dependent mechanics of collagen fibrils at the sub-fibrillar level remain poorly understood. This pioneering study employs atomic force microscopy (AFM) nano-rheometry and indentation testing to examine the viscoelastic characteristics of individual collagen type I fibrils at the ultrastructural level within distinct topographical zones, specifically focusing on gap and overlap regions. Our investigation has unveiled that collagen fibrils display a viscoelastic response that replicates the mechanical behavior of the tissue at the macroscale. Further, our findings suggest a distinct viscoelastic behavior between the gap and overlap regions, likely stemming from variances in molecular organization and cross-linking modalities within these specific sites. The results of our investigation provide unequivocal proof of the temporal dependence of mechanical properties and provides unique data to be compared to atomistic models, laying a foundation for refining the precision of macroscale models that strive to capture tissue viscoelasticity across varying length scales.

## Introduction

Collagens are the main structural proteins in the vertebrates and widely exist in both soft and hard tissues (1-3). According to the function and domain homology, collagens can be classified to fibril forming collagens, fibril-associated collagens with interrupted triple helices; network-forming collagens, transmembrane collagens, endostatin-producing collagens; anchoring fibrils, and beaded-filament collagens (4). Type I collagen monomer is the predominant molecule in fibrillar collagens found in nearly all connective tissues (5-7). Although type I collagen is the most abundant fibril-forming collagen, it assembles into heterotypic fibrils together with type III and type V collagens in tissues such as the lung, skin, and blood vessels, where type III collagen contributes to the formation of thinner fibrils (8, 9).

Fibrillar collagen constitutes the primary protein in bone, skin, tendons, ligaments, the sclera, cornea, and blood vessels (1, 2, 8-13). It provides the structural framework for these tissues, and its proper formation and maintenance are vital for their function (1, 2). It is crucial for bone extracellular matrix (ECM) mechanical competence and a key structural component of many other tissues (8, 11, 14). Moreover, fibrillar collagen through interaction with specific cell receptor including integrins regulates the functionality of cells in tissue (15-17). Therefore, understanding the mechanical behavior of collagen is essential for developing accurate models of biological systems such as the mechanics of blood flow through arteries, the deformation of cartilage under load(18) or the response of bone to dynamic loading (10, 19).

The remarkable mechanical attributes exhibited by collagen fibrils stem from their intricate hierarchical organization (Fig. A1), whose comprehension is crucial to deciphering its physiological and pathological relevance (20). At the macro-scale (i.e., whole tissues and collagen fibers), the mechanics of collagen-based tissues has been investigated over several decades. Fewer studies were reported at the micro and nanoscales (8) but it is widely recognized that collagen-rich tissue exhibits viscoelastic behavior, as evidenced by numerous creep and relaxation tests conducted on tendons and ligaments (21-23). Likewise, computational studies on collagen molecules and microfibrils revealed that the structural elements of collagen fibrils exhibit a viscoelastic response (24-26). This highlights the need for systematic experimental studies on the viscoelastic behavior of individual collagen fibrils. Various studies have explored the mechanical response of collagen fibrils using several advanced techniques(27). For example, MEMS-based stretching experiments have enabled precise control and measurement of mechanical forces, allowing for detailed analysis of fibril properties under various stretching conditions (28-33).

Shen et al. (31) investigated the viscoelastic properties of collagen fibrils isolated from sea cucumber dermis. Using a MEMS platform, *in vitro* coupled creep and stress relaxation tests were performed. They found that the relaxation time of single fibrils is significantly shorter than that of the source tissues, suggesting the influence of ground substance on tissue relaxation (31). Additionally, the elastic modulus was found higher in the first test compared to subsequent tests, indicating initial loading effects (31). Gautieri et al. (34) investigated collagen viscoelasticity at the molecular level and its relationship to connective tissue mechanics, addressing the challenges in linking molecular structure to mechanical properties. Their MEMS-based experiments, revealed that collagen fibrils can stretch up to 180% with a nominal strength of 0.5 GPa (34). Using atomistic modeling and comparing results with creep and relaxation tests of isolated fibrils, they found that individual collagen molecules exhibit non-linear viscoelastic behavior, while isolated fibrils fit the Maxwell-Weichert model (34). Additionally, X-ray diffraction has provided detailed insights into structural changes and mechanical behavior at the molecular level (22, 35-38). Likewise, AFM-based stretching has supplied crucial data on tensile strength and elasticity by pulling the fibrils and recording their force-extension behavior (39-42). For example, AFM bending techniques have assessed fibril flexibility and bending stiffness (43, 44). Additionally, AFM indentation has been employed to measure local mechanical properties, such as stiffness and hardness, by pressing a sharp tip into the fibril surface (6, 8, 9, 45-49). Using AFM indentation testing, Minary and Yu (50) investigated stiffness inhomogeneity within the substructural regions of collagen type I. Baldwin et al. investigated the structural and mechanical heterogeneity of collagen fibrils isolated from bovine tendon, demonstrating variations in elastic modulus along the length of the fibril (51-53). Gisbert et al investigates elastic modulus and loss tangent heterogeneity along collagen fibrils using high-speed bimodal AFM mapping (54). Despite these extensive studies, a thorough understanding of viscoelasticity found within individual collagen fibrils and their intricate substructures remains elusive.

On the modeling side, a number of constitutive and structural models have been developed to describe the viscoelastic behavior of collagen at the ECM fiber level and fibrillar levels (26, 37, 55-60). For instance, Silver et al. (61) examined strain rate dependence in self-assembled type I collagen’s stress-strain curves, and found that the elastic response is strain rate-independent, while the viscous response exhibits increased stress with strain rate. They observed that the elastic rate independent response matches the spring-like behavior of collagen’s flexible regions, and that collagen’s thixotropy results from fibrillar subunits slipping during tensile deformation (61). Sopakayang et al. (62) performed incremental stress relaxation tests on dry collagen from rat tail tendons and quantified viscoelastic properties, factoring in the contributions of microfibrils, interfibrillar matrix, and crosslinks. Their model, based on stress-strain and stress relaxation data, predicted that collagen fibrils relax faster with more crosslinks (62). Gautieri et al. (26) explored the viscoelastic properties of collagen molecules through atomistic modeling, focusing on the link between molecular structure and mechanical properties. Using the Kelvin-Voigt model, they assessed the impact of amino acid sequence on viscoelastic parameters, identified specific relaxation times for single collagen fibrils, and found that intramolecular hydrogen bonds significantly influence viscoelastic properties by affecting backbone rigidity and viscosity (26).

Deciphering the intrinsic viscoelastic properties of soft tissues has posed a longstanding challenge (8-10, 12, 63) so here we report an innovative approach based on AFM Nano-Dynamic Mechanical Analysis (Nano-DMA), often referred to as AFM Nano-rheometry, as well as direct AFM indentation to elucidate the local viscoelastic properties of collagen fibrils at the nanoscale with unprecedented precision (64-66). We summarize our results investigating the heterogeneous, time-dependent properties of single collagen fibrils at the ultrastructural level. We also correlate the experimental observations to the ultrastructural features of collagen.

## Materials and methods

### Fibril preparation

Human tropocollagen monomers (VitroCol) were sourced from Advanced BioMatrix Inc., Carlsbad, CA. The type I atelocollagen was supplied in acidic solutions with concentration of 3mg/mL, and at pH 2. To create the collagen fibrils, we first combined 20 μL of 200 mM disodium hydrophosphate (Na_2_HPO_4_, pH=7), 6 μL of sterile deionized water (ddH_2_O), and 10 μL of 400 mM potassium chloride (KCl) in an eppendorf centrifuge vial. We then added 4 μL of tropocollagen solution and mixed the solution at room temperature (Fig. 1). Subsequently, we placed the vial in a commercial cell culture incubator at 37°C for 8 hours. Upon completion, the solution had a slightly cloudy appearance and was primarily composed of native collagen type I fibrils(9). Lysyl oxidase was absent; therefore, enzymatic cross-linking density is negligible compared to that of native tissue.

**Figure 1:**
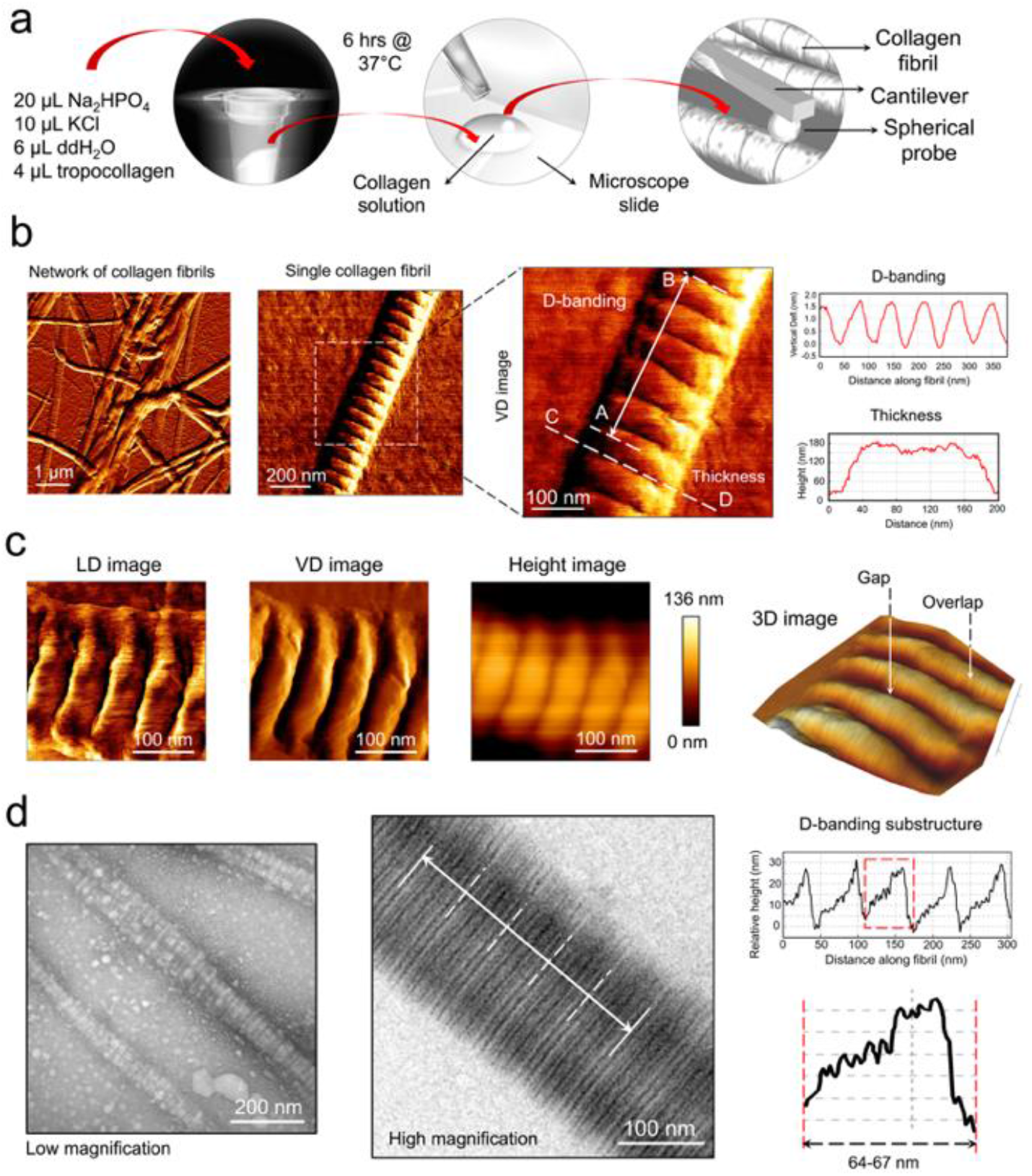
a: Illustration of the methodology employed to induce fibril formation *in vitro* for tropocollagen type I, followed by sample deposition on slide for AFM imaging in dry and mechanical testing in hydrated states(9). b: Vertical deflection (VD) AFM images of networks and isolated collagen fibrils with apparent D-banding periodicity in hydrated state. Fibril thickness and variation in the D-banding pattern on a representative single fibril, revealed by AFM image analysis are shown. c: Lateral deflection (LD), VD and height AFM images of a portion of an isolated collagen fibril in dehydrated state. In dry conditions, fibril height decreases due to dehydration, as shown in the bar graph of the height image in c. b & c exhibit the D-banding of ∼64-67 nm, with repetitive pattern of gap and overlap regions along the fibrils (8, 9). d: Transmission electron micrographs of fibrillar collagen showing the sub-regions of the D-banding pattern. A repetitive pattern of gap and overlap regions along collagen fibrils, as well as variations in this pattern on representative single fibrils, revealed by image analysis.

### Transmission electron microscopy (TEM)

For the TEM imaging of collagen fibrils, the solution was deposited on a glow-discharged, carbon-coated 200 mesh copper electron microscopy grid (Electron Microscopy Sciences, EMS, Hatfield, PA). After two minutes, excess solution was removed with filter paper, and the grid was left face-up for 30 minutes. The grid was washed with six drops of ddH2O for 10 minutes, fixed with 1% glutaraldehyde in ddH2O for two minutes, and washed again with six drops of ddH2O for 15 minutes (9). It was then dried at an angle on filter paper and counter-stained with 2% uranyl acetate for 30 minutes. After drying at an angle on filter paper, the grid was placed face-up on filter paper for 30 minutes. TEM imaging was conducted using a Tecnai 12 microscope (FEI Electron Optics) equipped with a tungsten filament at 120 kV and an AMT XR80C CCD Camera System at 50,000x magnification. The TEM images were analyzed using ImageJ software to measure fibril D-banding (Fig. 1).

### Atomic force microscopy

Indentation of collagen fibril subregions and entire fibrils at various speeds was performed using a JPK NanoWizard IV AFM (Bruker Nano, Berlin, Germany) mounted on an inverted epifluorescence Zeiss Axiovert 200M microscope (Carl Zeiss Microscopy, Göttingen, Germany). Dynamic mechanical characterization was subsequently conducted using a JPK NanoWizard PURE AFM (Bruker Nano, Berlin, Germany) on the same microscope platform. Reduced scan areas were selected to obtain the structural details of the fibrils. MSNL-10 silicon nitride cantilevers (Bruker, Mannheim, Germany) with a spring constant of 0.01--0.1 N/m and a nominal tip radius of ∼2 nm were used for high-quality imaging in an ambient environment of 20--25°C. Biosphere Au Reflex (CONT-Au) cantilevers (Nanotools USA LLC, Henderson, NV) with a nominal spring constant of 0.2 N/m, a length of 450 μm, and a nominal resonance frequency of 13 kHz in air were used. The cantilevers featured integrated spherical tips with radii of 20 nm (± 10%) and 50 nm (± 10%) (Fig. 2). The smaller radius tip was used for measurements within specific subregions, while the larger tip was applied for assessing the mechanical properties of the fibril as a whole. These cantilevers were employed for both imaging and subsequent indentation and Nano-DMA measurements. While some images were acquired in the dry state to clearly highlight the overlap and gap regions, those used for force measurements were captured in the hydrated state to ensure accurate selection of indentation and Nano-rheometry locations. All AFM mechanical measurements were performed in wet environment with fibrils being fully submerged in phosphate buffered saline (PBS) (Fig. 2). Before each test, the deflection sensitivity of the cantilever was calibrated by engaging the cantilever on a microscope slide in hydrated state. The precise spring constant of the cantilever was calibrated with the thermal noise fluctuations by fitting the first free resonance peak of the AFM cantilever to that of a simple harmonic oscillator using the JPK software (8). During the test, controlled deformation was applied to the sample in hydrated state and the compressive feedback forces were recorded and measured through cantilever deflection (Figs. 3 & 4).

**Figure 2:**
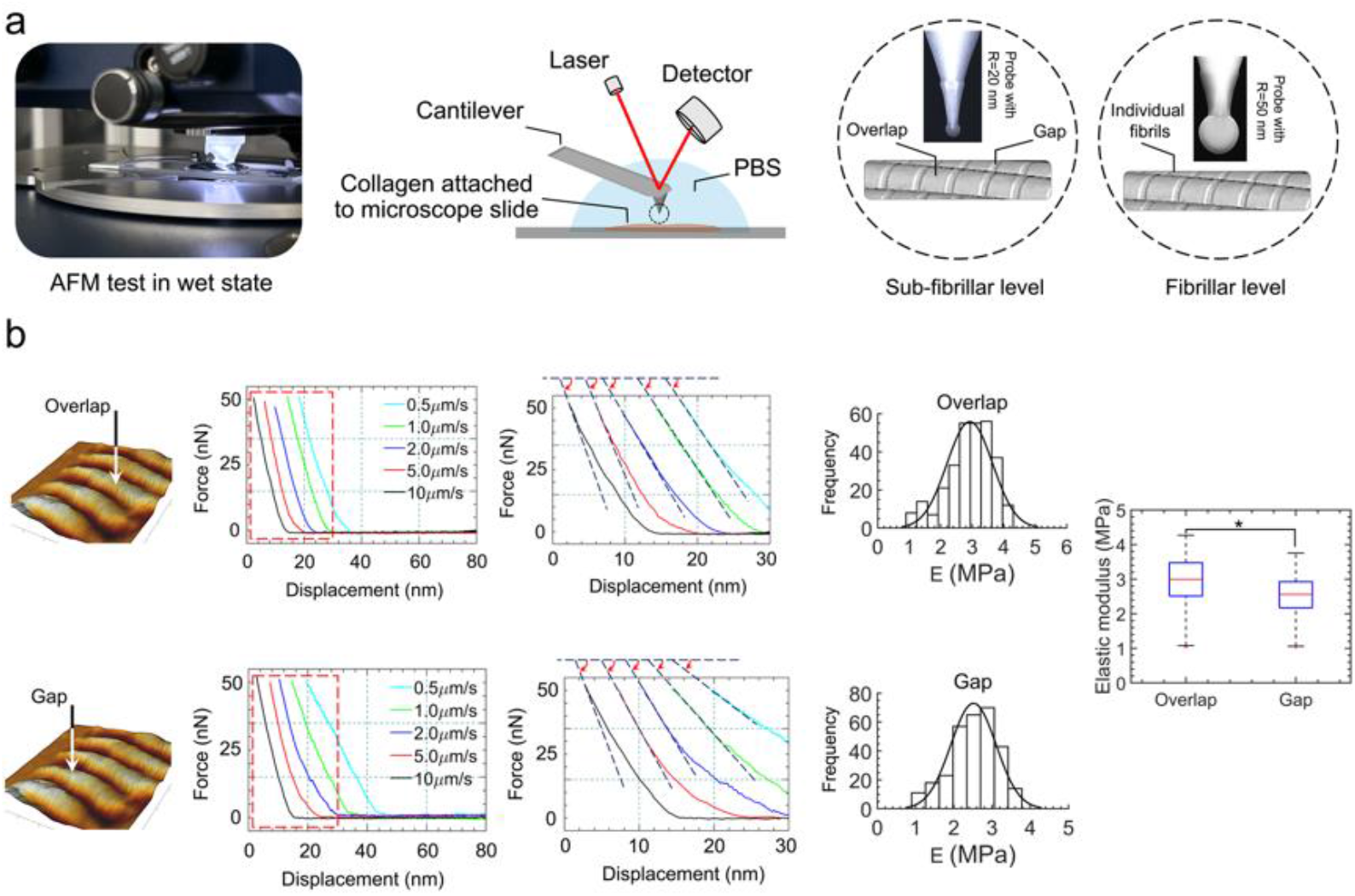
a: AFM testing at the sub-fibrillar and fibrillar levels in hydrated state. b: Force versus Z-displacement curves showing the relative increase in the slope of the initial portion of the unloading curve--indicative of elastic recovery--with increasing indentation speed in both the overlap and gap regions. Right panel in b: Elastic moduli of the overlap (2.92 ± 0.72 MPa) and gap (2.52 ± 0.59 MPa) regions at 10 μm/s, highlighting variations in stiffness with statistically significant differences (p<0.0001).

**Figure 3:**
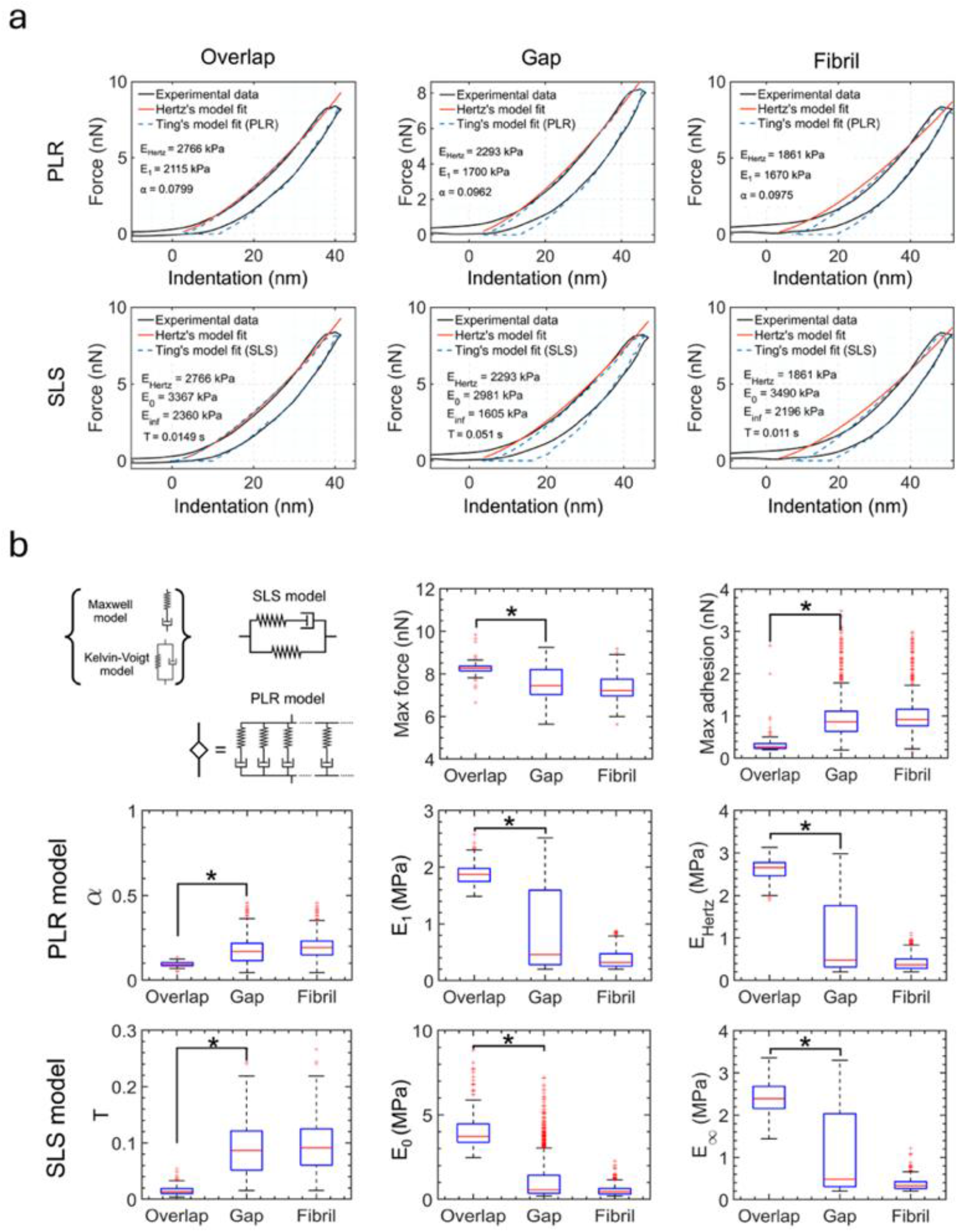
a: Representative indentation curves showing force against corrected tip displacement (i.e., Z-piezo displacement minus cantilever deflection) with fits from the PLR and SLS models, based on Ting’s solution and the Hertzian contact model, for the overlap and gap regions, as well as for the entire collagen fibril. AFM force-indentation curves are used to extract the viscoelastic parameters of type I collagen fibrils within these distinct sub-regions. b: SLS and PLR viscoelastic model parameters for overlap and gap regions and the entire fibril. The fitting parameters derived from both models reveal notable differences between the gap and overlap regions, as well as when analyzed across the whole fibril.

**Figure 4:**
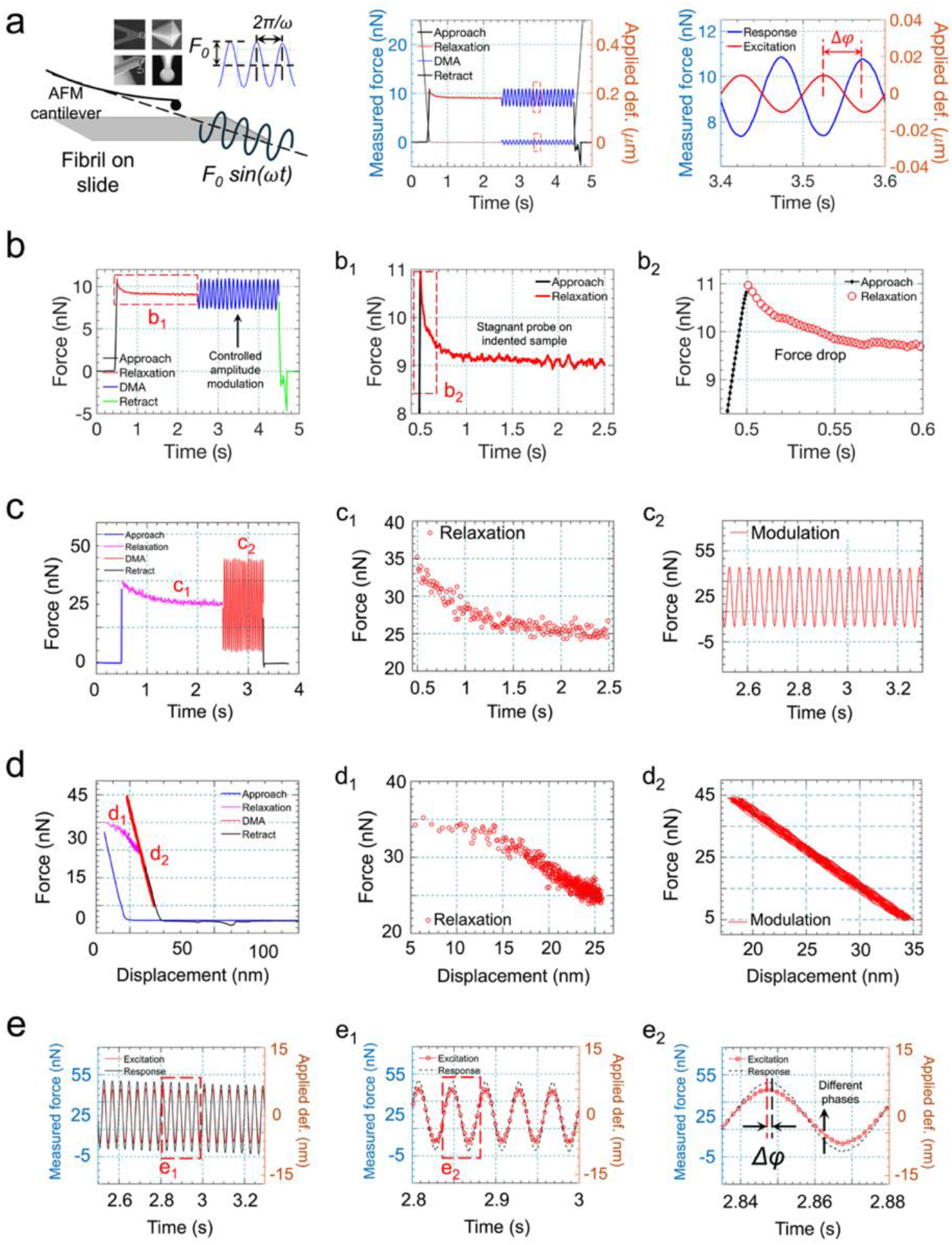
Nano-dynamic mechanical analysis (Nano-DMA) methodology employed to characterize the dynamic behavior of fibrillar collagen type 1. a: An AFM probe serves as both the force applicator and the displacement sensor. The tip applies a controlled oscillatory force on the sample’s surface while simultaneously measuring the material’s response. The storage and loss moduli are achieved through the analysis of the phase lag between the applied mechanical excitation and the resultant material response. The inset shows the SEM images of sharp and spherical AFM probes used in this study. b & c: Sample AFM curves for gap and overlap regions in time domain showing the rela xation and the modulation phases of the deformation because of the applied force. d) A force curve in a Nano-DMA experiment in the Z-displacement domain (i.e., the sum of tip displacement and cantilever deflection) that includes approaching and indenting with a maximum indentation depth of 10 nm, followed by a force-relaxation resulted from a pause in the indented state, and a sinusoidal response due to a harmonic excitation of 20 cycles in 2 sec prior to retracting. c_1 &_ d_1_: relaxation behavior after 2 sec in time and displacement domains for an overlap region. The net maximum indentation force *F*_*max*_ drops from ∼35 nN to ∼25 nN (∼40%) prior to approaching an asymptote at t=2.5 sec. c_2_ & d_2_: Recorded oscillating force response of 20 cycles in 2 sec in time and displacement domains for an overlap region. e: Simultaneous display of sinusoidal excitation and the harmonic response of the material with a slight phase lag Δ*φ* depicted in e_2_. The phase lag Δ*φ* between the excitation and the response yields the viscoelastic properties (i.e., the storage and loss moduli) of the sample.

### AFM direct viscoelastic characterization

The numerical processing of the F-Z curves was done using MATLAB scripts (The MathWorks, Natick, MA) developed in the previous works(63, 67) with the utilization of the Ting’s model(68):

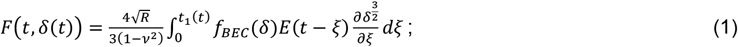

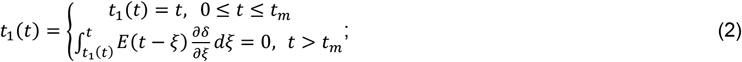

where F is the force acting on the cantilever tip; *δ* is the indentation depth; *t* is the time initiated at the contact; *t*_*m*_ is the duration of the approach phase; *t*_1_ is the auxiliary function determined by Eq. 2; *ξ* is the dummy time variable required for the integration; *v* is the Poisson’s ratio of the sample (assumed to be time-independent and equal to 0.5); R is the radius of the indenter; *f*_*BEC*_(*δ*) is the bottom-effect correction factor(69); and *E*(*t*) is the Young’s relaxation modulus for the selected rheology model.

Here we used two rheology model. The first model was the power-law rheology (PLR) model (Fig.3) (70, 71):

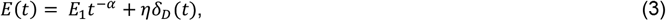

where *E*_1_ is the relaxation modulus at t = 1 s (scale factor of the relaxation modulus); *α* is the power-law exponent; *η* is the Newtonian viscous term (with Pa*s units); and *δ*_*D*_(*t*) is the Dirac delta function. A larger *α* value means a larger amount of relaxation; materials exhibit a solid-like behavior at *α* = 0, and a fluid-like behavior at *α* = 1. The PLR model described by Eq. (3) was successfully used for the description of cell mechanics in several previous studies(70, 72-74). The second model was the standard linear solid (SLS) model with the following equation for the Young’s relaxation modulus:

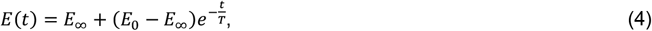

where the three parameters are: *E*_0_ is the instantaneous modulus, *E*_∞_ is the infinite (long-term, equilibrium) modulus and T is the relaxation time of the material. The Young’s modulus with the assumptions of the Hertz’s theory, YM (“apparent” elastic modulus), was also calculated from the approach part of the force curves(75).

### Nano-rheometry or Nano-Dynamic Mechanical Analysis (Nano-DMA)

The foundation of the nano-DMA technique is rooted in the presumption that the contact area between the AFM tip and the sample remains consistent throughout the oscillation of the tip. This presumption finds support in the substantial adhesive forces inherent to soft materials. To grasp this presumption, consider a spherical probe with a tip radius denoted as R, dynamically penetrating an adhesive viscoelastic substance. The indentation process is characterized by sinusoidal waveforms. Consider indentation *δ* and force *F* of the indenter in contact with a viscoelastic medium in an equilibrium state (*δ*_0_,*F*_0_). Small-amplitude oscillations with angular velocity ω around that state are described as

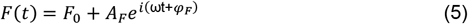

and

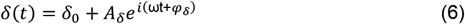

The amplitudes *A*_*F*_ and *A*_*δ*_ and the phase difference Δ*φ* = *φ*_*F*_ − *φ*_*δ*_ can be measured in an experiment. As the material’s displacement (or its response) lags behind the applied sinusoidal force, the material modulus adopts a complex nature, denoted as *E*^*^. The stiffness of the material, expressed in terms of the force and displacement amplitudes as *F*_0_/*δ*_0_, is intricately linked to the complex modulus as

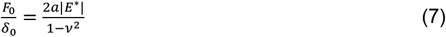

with *a* being the contact radius between the sample and the spherical probe, and *v* denoting the Poisson’s ratio. The force is alternatively expressed as

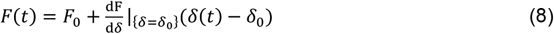

with

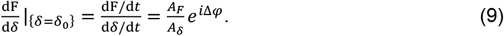

Upon substitution of *δ*(*t*),

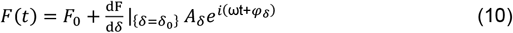

The complex elastic modulus *E*^*^ with a spherical indenter of radius R, according to Sneddon’s (1965) elasticity model, is

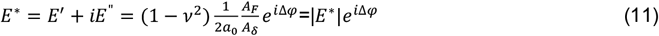

with the storage modulus *E*^′^ and the loss modulus *E*^”^ being the real and imaginary parts of *E*^*^, viz.,

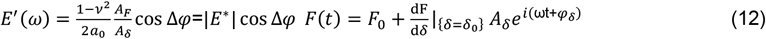

and

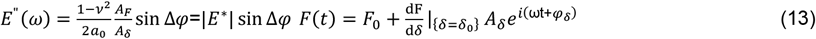

The loss tangent,

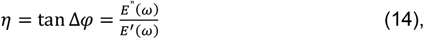

is an indicator of solid-like (tan Δ*φ* ≪1) or liquid-like (tan Δ*φ* ≫1) behavior.

### Data collection and statistical analysis

After imaging the fibrils on each slide in hydrated state, the AFM mode was switched from imaging to force spectroscopy. For each fibril, multiple locations were carefully selected within both the gap and overlap subregions, with indentation points recorded separately to ensure that data from each region remained distinct (See Supplementary Information). Indentation tests were performed at the selected sites, and corresponding raw force curves were collected. The same procedure was applied for Nano-rheometry measurements. Although the D-banding appeared less pronounced in the hydrated state, the indentation points were selected under hydrated conditions (see Supplementary Information). Both indentation and Nano-rheometry tests were conducted using four biological replicates. For each replicate, 10 to 20 fibrils were analyzed across three slides, with a minimum of ten gap/overlap region pairs tested per fibril. At each selected site, indentations were repeated to confirm elastic deformation and no permanent deformation on the fibril (See Supplementary Information). Raw force-distance curves and Nano-rheological data were obtained, with key parameters (e.g., approach/retract speed kept consistent across all measurements.

AFM data analysis was performed using MATLAB (version 2018) and JPK Data Processing software (version 6.3). Data visualization and statistical analysis for Figures 5 and 6 were carried out in GraphPad Prism 10. Statistical differences were assessed using one-way ANOVA with a significance level of 0.05. Data are presented as mean ± standard deviation (SD) unless otherwise noted.

**Figure 5:**
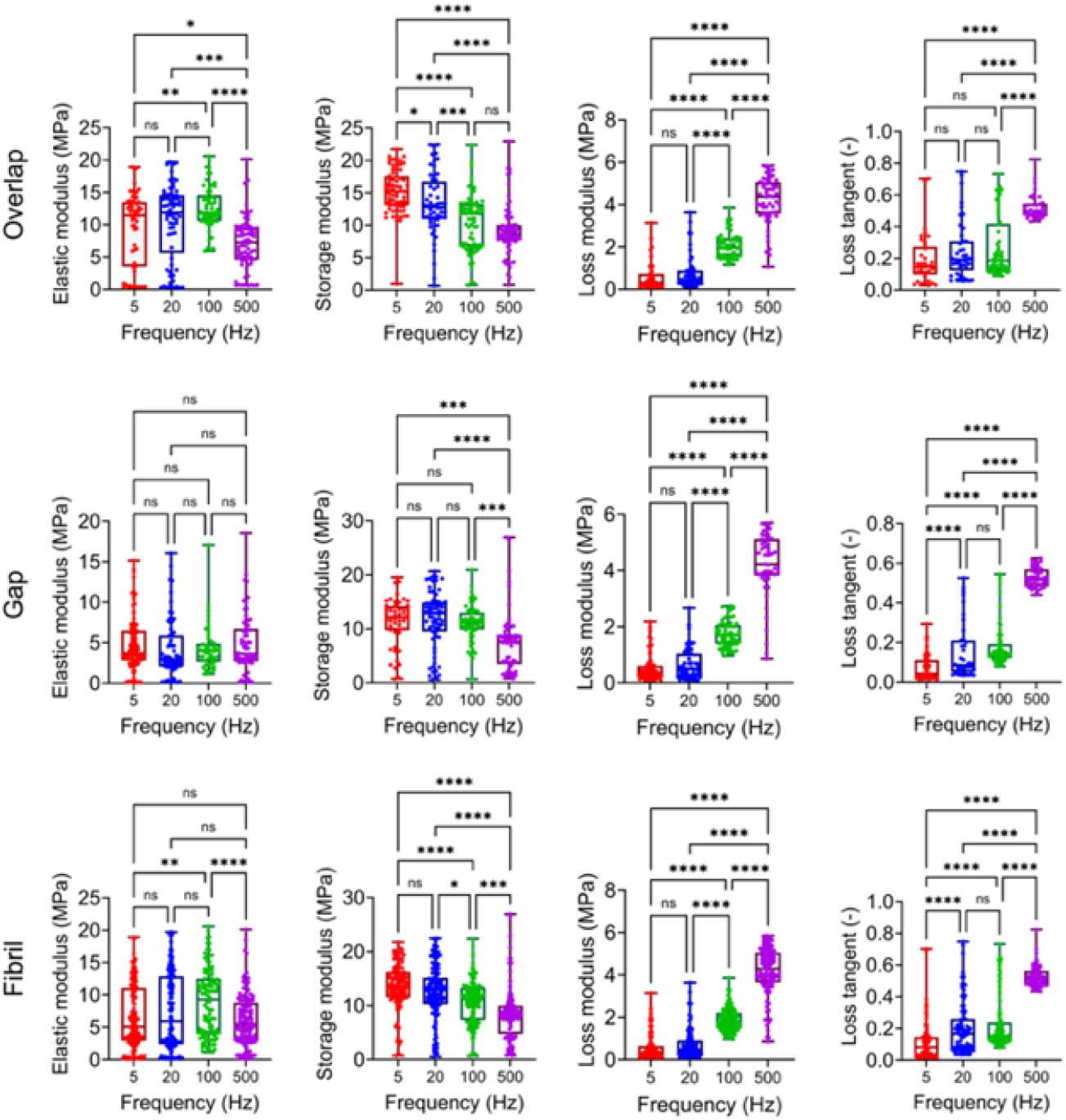
Graphs of elastic modulus *E*, storage modulus *E*^′^, loss modulus *E*^”^, and loss tangent *η* measured for overlap and gap regions at 5 Hz (red), 20 Hz (blue), 100 Hz (green), and 500 Hz (purple), as well as the whole fibril at several frequencies at the cantilever oscillation amplitudes of 5 nm.

**Figure 6:**
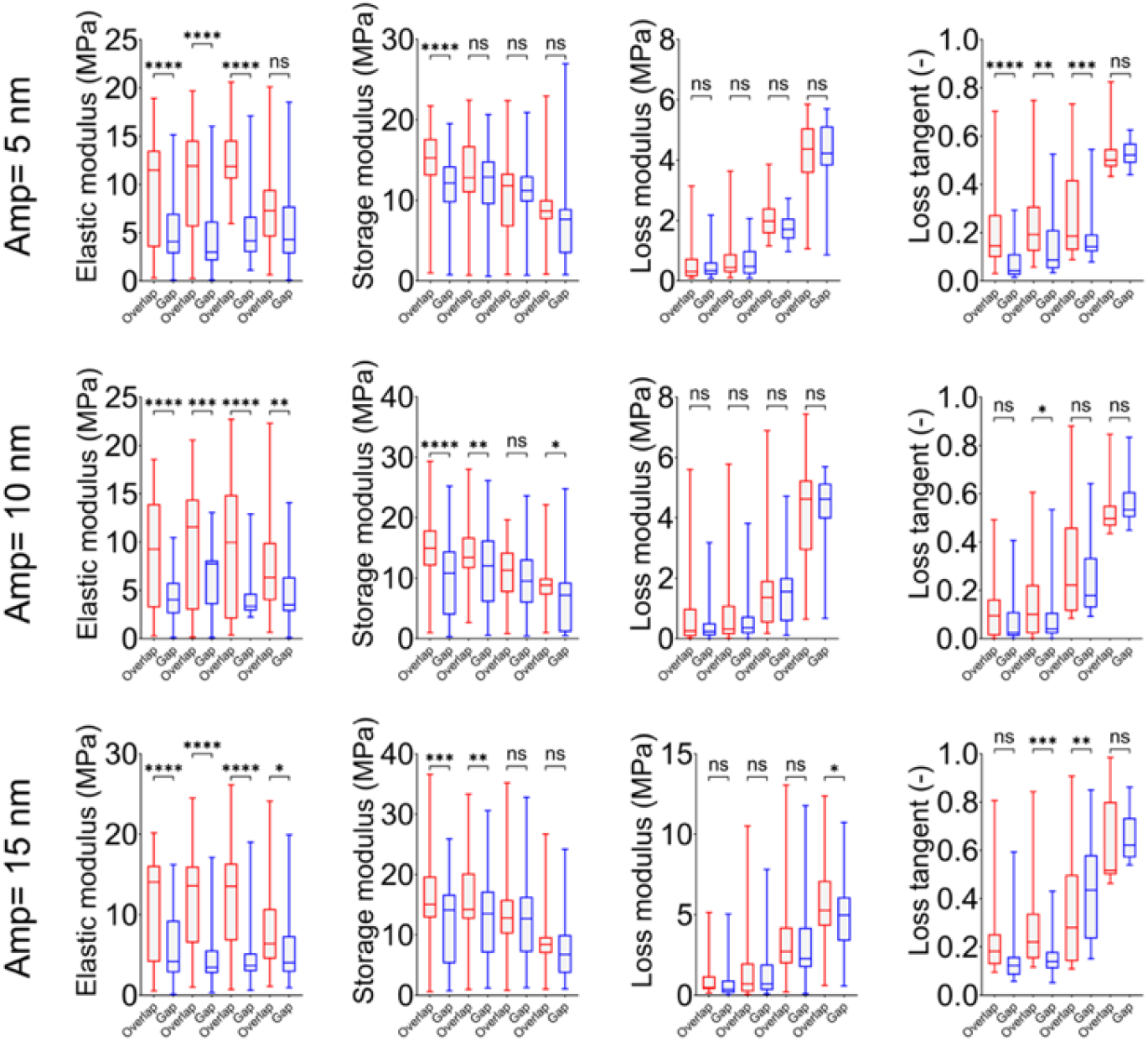
Comparison of elastic modulus *E*, storage modulus *E*^′^, loss modulus *E*^”^, and loss tangent *η* between overlap (red) and gap (blue) regions, at 5 Hz (first pair of data columns in every plot), 20 Hz (second pair), 100 Hz (third pair), and 500 Hz (last pair) at three selected cantilever oscillation amplitudes of 5, 10 and 15 nm. The elastic modulus in the overlap region is consistently observed to be greater than that of the gap region, highlighting a distinct mechanical disparity between the two domains. In the majority of cases, the loss modulus in the overlap region is significantly higher than in the gap region, indicating increased energy dissipation within the overlap domain.

## Results and discussion

### Viscoelasticity characterization directly from indentation testing

Indentation testing at varying speeds demonstrated an increase in the elastic modulus of collagen fibrils in both the overlap and gap regions (Fig. 2), aligning with prior *in-vitro* and *in-vivo* observations (8, 9). Elastic moduli of the overlap (2.92 ± 0.72 MPa) and gap (2.52 ± 0.59 MPa) regions at indentation speed of 10 μm/s showed statistically significant differences (p<0.0001) (Fig. 2b). To rigorously assess the time-dependent mechanical behavior, we applied the approach developed by Efremov et al.(63), which employs a relaxation-based Hertzian model to directly extract viscoelastic properties from the indentation data. Significant spatial variations in mechanical behavior were observed between the overlap and gap zones (p<0.0001). As illustrated in Figure 3a, maximum adhesion force varied significantly, with the overlap region showing the lowest adhesion force (0.34 ± 0.03 nN) and the gap region exhibiting a substantially higher value (0.92 ± 0.02 nN). Correspondingly, the maximum force also differed across these regions (Fig. 3b), reaching its highest value in the overlap region (8.26 ± 0.02 nN) and declining in the gap region (7.58 ± 0.02 nN). These trends suggest region-dependent mechanical interactions, possibly related to local molecular organization and crosslinking density.

Analysis using the Power-Law Rheology (PLR) model revealed distinct viscoelastic characteristics across fibril regions (Figure 3b). The viscoelastic parameter α was lowest in the overlap region (0.09 ± 0.001), while the gap region exhibited a significantly higher value (0.19 ± 0.002), indicating a more pronounced time-dependent response in the gap region. Similarly, the instantaneous elastic modulus (*E*_1_) showed marked spatial variation, with the highest value obtained in the overlap region (1.87 ± 0.01 MPa), decreasing significantly in the gap region (0.82 ± 0.02 MPa). The Hertz constant also followed this trend, reaching a maximum in the overlap region (2.6 ± 0.02 MPa) and a minimum in the gap region (0.95 ± 0.03 MPa). This consistent reduction in elastic modulus across the gap region suggests a localized stiffness reduction potentially associated with lower fibrillar density or reduced molecular alignment (Fig. A1).

The Standard Linear Solid (SLS) model supported these observations, highlighting pronounced regional differences in viscoelastic properties (Figure 3b). The *E*_*0*_ modulus in the SLS model was highest in the overlap region (4.15 ± 0.08 MPa) and lowest in the gap region (1.25 ± 0.04 MPa), analogous to the trend observed in the PLR model. Similarly, the *E*_∞_ modulus was significantly greater in the overlap region (2.4 ± 0.02 MPa) compared to the gap region (1.06 ± 0.04 MPa). The constant *T* also varied substantially, with a faster relaxation (lower *T* value) in the overlap region (0.02 ± 0.001) and slower relaxation in the gap region (0.09 ± 0.001). These findings further support the hypothesis that mechanical and viscoelastic properties of collagen fibrils are spatially heterogeneous, with the overlap regions exhibiting greater stiffness and reduced time-dependent deformation relative to the gap regions. These observations may stem from collagen’s molecular rearrangement (Fig. A1) and sliding during loading, which lead to hysteresis. Irreversible deformation limits energy recovery during unloading, resulting in energy dissipation and contributing to the material’s viscoelastic behavior.

### Viscoelasticity characterization from Nano-Rheometry

Dynamic nanomechanical properties are recognized as crucial drivers of cell biology (76). However, due to technological challenges, these properties have not been studied at multiple scales in the context of tissues. Additionally, the development of viscoelastic properties in scaffolds and their evolution during tissue formation remain poorly understood. We addressed this technological barrier by employing AFM Nano-rheological characterization, which enables the measurement of dynamic mechanical properties (storage and loss moduli) across a wide range of sample scales and frequencies (Fig. 4). Using this method, we measured the dynamic moduli of collagen type I fibrils, as the fundamental building blocks of the tissue scaffold. This approach enables the characterization of a nano-rheological signature of fibril nano-mechanics, particularly in diseased tissues, which are characterized by increased stiffness, decreased rate-dependent stiffening, and reduced energy dissipation. Further, by testing the sub-fibrillar regions of collagen type I, we revealed a stepwise emergence of rheological properties as a function of fibril hierarchy and scale.

In Nano-DMA measurements, excitation and response signals are shown on separate axes (Fig. 4). The left axis, labeled “applied deformation”, reflects the designed loading set in the *Advanced Force Setting* module, which includes a sequence of indentation, pause, modulation, and retraction. During indentation, a static force deforms the sample. The probe is then held at the maximum depth to allow viscoelastic relaxation, followed by a sinusoidal modulation where the probe oscillates while remaining in contact with the sample in the indented state. Finally, the probe is retracted to complete the cycle. This sequence is illustrated in the Figure 4a, showing the applied mechanical stimulus and the resulting deformation. The sample’s response is recorded in terms of forces calculated using cantilever deflection and the calibrated stiffness of the cantilever. Initially, force increases with indentation, then gradually decreases during the pause as the sample experiences force relaxation (Figs. 4b_1,_ c_1 &_ d_1_). During the modulation phase (Fig. 4e), the sample exhibits sinusoidal deformation with a phase lag relative to the excitation (Fig. 4e_2_), indicating viscoelastic behavior. The force then returns to baseline upon retraction.

As shown in Figure 4, an AFM probe, with both sharp and spherical tips visible in SEM images, functions as the force applicator and displacement sensor on sub-fibrillar regions. The tip applies a controlled oscillatory force to the sample’s surface while measuring the material’s response. Storage and loss moduli are derived from the phase lag between the applied force and the material’s response. AFM curves for gap and overlap regions show relaxation and modulation phases due to the applied force (Fig. 4b & 4c). A force-distance curve in a Nano-DMA experiment illustrates the process, including approach, indentation, force relaxation, and a sinusoidal response due to harmonic excitation of 20 cycles in 2 seconds prior to retraction in the overlap region (Fig. 4d). The relaxation behavior after 2 seconds shows the maximum indentation force *F*_*max*_ dropping from ∼35 nN to ∼25 nN (∼40%) before stabilizing at 2.5 seconds. An oscillating force response over 20 cycles in 2 seconds is shown in Figure 4c_2_. The indentation loop indicates a maximum depth of 10 nm in the overlap region, with relaxation followed by 20 cyclic loadings in the displacement domain shown in Figure 4d. Simultaneous sinusoidal excitation and harmonic response with a slight phase lag Δ*φ* are depicted in Figure 4e, illustrating the methodology used to characterize the dynamic behavior. The phase lag Δ*φ* between excitation and response was used to compute the storage and loss moduli. Conversely, as illustrated in Figure B1 (Appendix B), Nano-DMA testing on a glass slide, used as a rigid surface, demonstrates no relaxation in time or displacement fields, with zero phase difference between excitation and surface response.

As shown in Figure 5, the elastic modulus *E* of the collagen fibril, in the gap region, showed no significant change with increasing oscillation rate. Yet, this parameter exhibited a decrease, within the overlap region of the collagen fibril, from 9.45 ± 0.75 MPa to 7.13 ± 0.47 MPa (p<0.03). Within the gap region, the loss modulus *E”* surged from 0.51 ± 0.05 MPa to 4.2 ± 0.16 MPa (p<0.0001), whereas the storage modulus *E’* decreased from 11.32 ± 0.62 MPa to 7.59 ± 0.72 MPa (p<0.0003). A comparable alteration in *E’* and *E”* occurred within the overlap of the collagen fibril as the oscillation rate of the cantilever on the fibril increased (Fig. 5). The increase in *E”* and decrease in *E’* with oscillation rate may be partially attributed to changes in hydration levels affecting the fibril’s viscoelastic behavior (46, 77). TEM studies indicate that minerals preferentially nucleate in the less ordered gap regions of bone collagen (78, 79). This observation suggests a structural distinction between the gap and overlap regions. This is consistent with our findings on the viscoelastic properties, which show significant changes in the loss modulus *E”* in the gap region with increasing oscillation rates. This behavior suggests that the gap region may be more adept at energy dissipation, likely due to its structural irregularities and mineral content, making it more adaptable to varying mechanical stresses.

The reduction in elastic modulus with hydration is likely attributed to the formation of water bridges between peptide chains (46). This observation mirrors the mechanical behavior of the tissue when examined on a macroscopic scale, highlighting the differences between its properties in dry versus hydrated states (10). As depicted in Figure 6, when the material is examined under an oscillation with a 5 nm amplitude, the elastic modulus *E* in the gap region remained independent of the oscillation frequency. The elastic modulus was also lower than that in the overlap region (p<0.0001). Furthermore, under the same oscillation amplitude, the storage moduli in the gap and overlap regions were observed to differ (15.30 ± 0.41 MPa and 11.32 ± 0.62 MPa respectively; p<0.0001) (see Fig. 6). However, neither of them underwent significant changes with increasing frequency (Fig. 6). Concerning the loss modulus, no alterations were detected within either the overlap or gap regions as the frequency increased (Fig. 6). When the cantilever oscillation amplitude was increased to 10 nm, the elastic modulus *E* in the gap region was observed to be lower than that in the overlap region (p<0.0001). X-ray diffraction studies indicate that the overlap region exhibits roughly twice the order of the gap region (80). This finding aligns with our observation that the elastic modulus *E* decreases with increasing oscillation rate in the overlap region, while it remains constant in the gap region. The higher degree of order in the overlap region likely makes it more sensitive to mechanical changes under dynamic conditions, whereas the more disordered structure of the gap region may enhance its mechanical stability across varying oscillation rates (81).

The storage modulus *E’* in the gap region was also lower than that in the overlap region at both 5 Hz and 500 Hz, representing the lowest and highest applied frequencies, respectively (p<0.0001 & p=0.035, respectively) (Fig. 6). As the oscillation amplitude of the cantilever increased to 15 nm, the elastic modulus *E* in the gap region consistently stayed lower than that in the overlap region across varying frequencies (p<0.0001). The storage moduli differed significantly between the gap and overlap regions at 5 Hz (16.19 ± 0.93 MPa and 11.9 ± 0.83 MPa, p<0.0005), but this trend did not persist with increasing frequency (Fig. 6). At a frequency of 500 Hz, the loss modulus in the gap region was observed to be lower compared to that in the overlap region (5.69 ± 0.25 MPa and 4.8 ± 0.28 MPa, p<0.047). The reduction in storage modulus *E’* observed in both regions with increasing oscillation rates highlights the viscoelastic nature of collagen fibrils. However, the differing behaviors in the gap and overlap regions emphasize the impact of structural heterogeneity on energy storage and dissipation. The gap regions, with their greater flexibility, exhibit higher energy dissipation (indicated by a higher *E”*) and show less dependence on oscillation rate for maintaining elastic properties. In contrast, the more ordered overlap regions display more significant changes in both *E’* and *E”* under dynamic loading. Due to the absence of the LOX enzyme in the in vitro model, the density of enzymatic covalent cross-links is expected to be negligible compared to mature tissues such as tendon and fibrotic scar (Fig. A1). As a result, the observed viscoelastic response is primarily governed by intrinsic molecular packing interactions, such as hydrogen bonding (82). Previous investigations have provided a comprehensive evaluation of the nanomechanical heterogeneity within the gap and overlap regions of individual type I collagen fibrils, revealing pronounced differences in their elastic and energy dissipation behaviors (50).

Continuous dynamic mapping, alongside static indentation, identified significantly higher energy dissipation in the overlap region compared to the gap region, pointing to distinct mechanisms of energy dissipation during dynamic deformation (50). Our dynamic mechanical analysis offers a precise quantitative characterization of these mechanisms, which likely involve molecular sliding and friction within the collagen matrix, driven by weak electrostatic and hydrophobic interactions. These interactions facilitate bond breaking and reformation under oscillatory loading conditions, underscoring the intricate ultrastructural mechanical complexity of collagen fibrils.

The loss tangent *η* serves as a comparative measure between energy dissipation and energy storage within a material. In both the overlap and gap regions, *η* values are consistently below 1 (Fig. 6), indicating that energy storage surpasses dissipation. This suggests that both regions predominantly exhibit elastic behavior, efficiently storing mechanical energy during deformation and recovering it upon unloading with minimal energy loss.

Nanoscale heterogeneity between gap and overlap regions in collagen can significantly influence tissue-scale function in pathological conditions like fibrosis (83) or aging (84). These variations affect local mechanical properties, molecular accessibility, and remodeling behavior, potentially leading to altered tissue stiffness or impaired cell-matrix interactions. In fibrosis, exaggerated cross-linking and altered nanostructure can stiffen tissues, disrupting normal function (83). In aging, changes in fibril organization may reduce elasticity and repair capacity (84). Understanding this heterogeneity is also critical for scaffold design, where replicating native nano-architecture can improve integration and mechanical performance.

We note that the inherent curvature of the D-banding pattern, particularly subtle differences in surface convexity or concavity between gap and overlap regions, may influence the precise tip-sample contact geometry, potentially contributing to local variability in stiffness and adhesion measurements. However, this effect is likely minimal given the small AFM tip radius and low applied loads used in our experiments.

### Molecular structures and associated deformation mechanisms

The augmentation of structural integrity and load-bearing capability in the fibril is a direct consequence of covalent cross-links formed between collagen molecules, meaning the enzymatically formed interfibrillar bonds including reducible, carbonyl-derived cross-links (85). These interfibrillar cross-links not only serve to stabilize the fibrillar architecture but also act as steadfast barriers, preventing any potential slippage between the constituent molecules. Moreover, the reiterated sequences of amino acids present in collagen molecules (Fig. A1), encompassing glycine, proline, and hydroxyproline, have the potential to foster the creation of hydrogen bonds linking neighboring α-polypeptide chains of collagen molecules. These hydrogen bonds play a pivotal role in fortifying the fibril’s stability and facilitating the transmission of forces along its elongated axis (86, 87). Furthermore, the quasi-crystalline molecular configuration within a collagen fibril engenders a proficient mechanism for load distribution, ensuring a uniform dispersion of forces across the fibril (88). The disparities in diameter along the fibril’s extent exert a pronounced influence on its mechanisms of deformation, thereby augmenting the broader spectrum of load distribution within the fibril. Robust segments of the fibril, endowed with greater thickness, manifest an enhanced capacity to withstand formidable loads, whereas more slender sections exhibit a greater tendency to be flexible. Water molecules within the fibril’s structure significantly contribute to load transfer along a single fibril through the hydration of tropocollagen molecules, facilitating the formation of hydrogen bonds between them (89). Moreover, the water-filled voids within the fibril confer heightened flexibility and enhanced resistance against compressive forces, such as those arising from indentation. These mechanisms synergistically contribute to impart remarkable strength to collagen fibrils. However, the contribution of each mechanism is contingent upon the distinct source tissue and prevailing physiological conditions (87). AFM studies reveal that the overlap region has a larger cross-sectional area and is less prone to unfolding compared to the gap region (51-54). Our results align with this observation, as we found a decrease in both the elastic modulus *E* and storage modulus *E’* in the overlap region. This decrease likely indicates a more compact and less flexible structure under mechanical stress. Conversely, the gap region exhibits higher strain resilience and an increased loss modulus *E”*, suggesting that it can accommodate greater mechanical deformation while maintaining its structural integrity, which corresponds to its greater tendency to unfold.

Hydration significantly influences the viscoelastic properties of collagen fibrils by modulating their molecular mobility and intermolecular interactions (45). In the hydrated state, water molecules infiltrate the collagen structure, disrupting hydrogen bonding and facilitating increased sliding between collagen triple helices and fibrillar subunits (45). This leads to a marked reduction in stiffness (lower storage modulus) and an increase in energy dissipation (higher loss modulus), reflecting a more compliant and viscous mechanical response.

Proline-rich collagen triple helices demonstrate higher elastic modulus than their hydroxyproline-rich counterparts, highlighting the critical influence of amino acid sequence on collagen mechanics (90). Supporting this, molecular dynamics “virtual creep” simulations show a time-dependent reorganization of hydrogen bonds under sustained load: hydroxyproline-rich motifs gradually lose intramolecular hydrogen bonds while forming new protein-water interactions, whereas proline-rich regions more stably preserve their intramolecular bonding (90). These findings suggest that sequence not only dictates triple-helix stability but also modulates its hydration state under mechanical strain. The load-induced increase in protein–solvent hydrogen bonding may expose polar groups, enabling the formation of new intermolecular hydrogen bonds, either direct or water-bridged, with adjacent collagen molecules or fibrils. This provides a plausible mechanism for stress-mediated coupling within the fibrillar network.

## Concluding remarks

Collagen, as the preeminent protein of the human body, assumes a pivotal role in sustaining the structural integrity and mechanical robustness of various tissues and organs. It ensures their functionality and resilience under numerous physiological processes and biomechanical stresses. The hierarchical organization of a collagen fibril (Fig. A1) furnishes it with exceptional mechanical properties, which are fundamental in comprehending its physiological and pathological responses. Particularly significant for biological tissues that undergo incessant loading and unloading cycles are the viscoelastic properties that describe a material’s capacity to deform and recover over time. Here we utilized AFM Nano-rheometry also called Nano-DMA, as well as direct indentation testing to unravel the ultrastructural viscoelastic attributes of ultrastructural subregions of collagen type I at the nanoscale. Our observations reveal that collagen fibrils exhibit viscoelasticity variations along the fibril, particularly within the overlap and gap subregions of the D-banding pattern. This variability may stem from the distinct molecular arrangements present in these subregions. The time-dependent properties of collagen fibrils, as elucidated by Nano-DMA, are markedly influenced by the inherent structural disparities between the gap and overlap regions. The gap region, characterized by its less organized and more pliable nature, demonstrates distinct mechanical responses in contrast to the more structured and rigid overlap region. These variations are reflected in the elastic, storage and loss moduli across a spectrum of amplitudes and frequencies, underscoring the intricate interplay between structural organization and mechanical behavior in a collagen fibril. Our results time-dependent properties at the nanoscale, which could be used in comparison with atomistic models, establishing a groundwork for refining the precision of macroscale models used for soft tissue mechanics across a range of length scales.

## Notes

The authors declare no competing financial interest.

## Acknowledgments

MA (PI) appreciates the faculty startup package from the University of South Florida, the technical support from Nanotools USA, and the Facility for Electron Microscopy Research (FEMR) at McGill University for providing access to Transmission Electron Microscopes.

## Appendices Appendix A: Collagen I synthesis and structural characterization

At the molecular level, collagen 1 is composed of a heterotypic triple α-helix derived from Col1 α1 and Col1 α2 chains typically in a 2:1 stoichiometric ratio (91). Each α-polypeptide chain of collagen consists of 1014 amino acids (92). The amino acids have been assembled with repeated permutation along each chain followed by the tripeptide sequence of an N-terminus, a series of Gly (glycine)-X-Y repeats, and a C-terminus. X and Y can be any amino acid except Gly, but are usually proline and hydroxyproline, respectively (93). α-chains form left-handed helices with 3.3 residues per turn and a pitch of 0.94 nm, compared with the common peptides, which form right-handed α-helices with a pitch of 0.54 nm and 3.6 residues per turn, suggesting that the α-chain of collagen is stretched but narrowly twisted (94, 95).

All three α-polypeptide chains of collagen undergo post-translational processing that are essential for the mechanical competence of collagen fibrils (96, 97). The lysine and proline residues get additional hydroxyl groups added to them via hydroxylase enzymes which require vitamin C, oxygen, and iron as cofactors (96, 97). Glycosylation of the selected hydroxyl groups on lysine with galactose and glucose is catalyzed by galactosyl transferase and glucosyl transferase (96, 97). Concomitant with hydroxylation and glycosylation modifications, the α-chain polypeptides of collagen initiate folding into a triple helix by registration and disulfide bonding of the carboxy-terminal ends of two α1(I) polypeptides and one α2(I) polypeptide to form the tertiary structure of pro-collagen (98, 99). Inside the triple helix, every third amino acid residue is located around the central axis, where the interspace along α-chains is the narrowest (100). Therefore, the glycine residues are suitable for the third amino acid residues because of its minimum volume (100). Besides, the free amino acid residues in the side chain extend outward perpendicular to the spiral axis and assist in formation of multiple hydrogen bonds in the spiral chain, which is essential for maintaining the stability of the helical structure (100) (Fig. A1).

In ECM, the telopeptides in C and N terminals consisting amino acid sequences are cleaved by metalloproteinases, forming tropocollagen (86). In the ECM, tropocollagen molecules assemble into a higher-order structure known as a collagen fibril, i.e., the fundamental unit of collagen fibers. The introduction of covalent cross-links between tropocollagen molecules aligns them end-to-end and parallel. Cross-linking is mostly initiated by members of the lysyl oxidase (LOX) family of copper-dependent amine oxidases (101, 102). The effect of LOX activity on fibrillar collagens is to convert lysine or hydroxylysine residues in the N- and C-terminal telopeptide regions to corresponding peptidyl aldehydes. Once formed, aldehydes spontaneously condense with other aldehydes or unreacted lysines and hydroxylysines to form a variety of intra- and intermolecular covalent cross-link (103). The precise arrangement of collagen molecules within the fibril structure creates a unique characteristic “D-banding” or “D-spacing” pattern that plays a critical role in determining the mechanical properties of the fibril (2, 80) (Fig. 1 & A1). The structure of an isolated fibril is continuous, and the intrafibrillar bonds are thus essential to guarantee the transfer of mechanical loads along the fibril. The load propagation mechanisms embedded within the structural framework of an individual collagen fibril can amalgamate a synergy of molecular interactions and architectural configuration. Collagen fibrils are strengthened through covalent intermolecular cross-links formed enzymatically during the assembly between the collagen molecules. The formation of intermolecular covalent cross-links in collagen is important in enhancing the mechanical stability of the collagenous tissues (20, 104).

**Figure A1:**
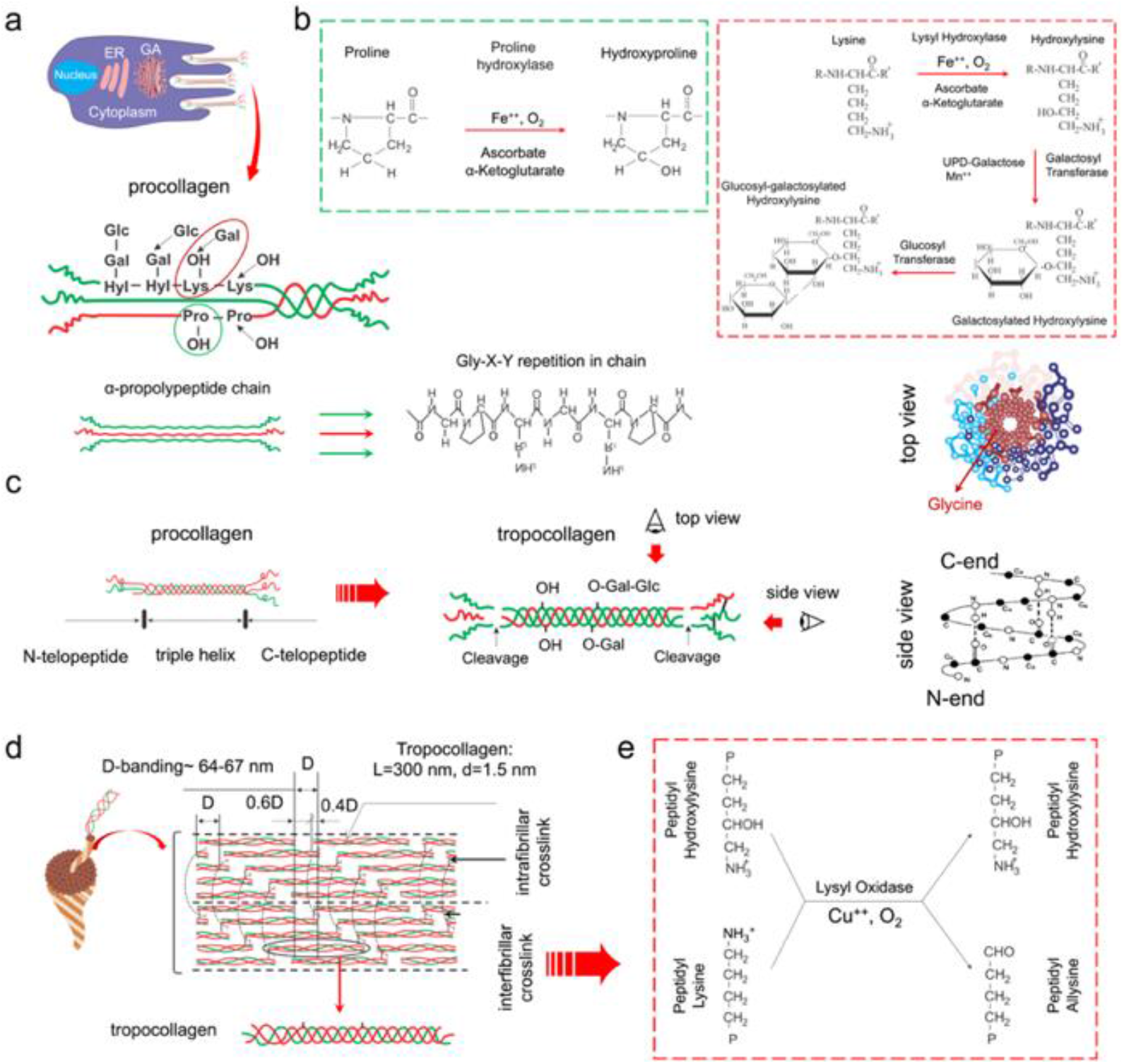
Collagen expression, synthesis, and posttranslational modification. a: Collagen, the most abundant protein in the human body, is encoded by specific genes in the cell’s nucleus. The α-pro-polypeptide chain has a tripeptide sequence of an N-terminus, a series of Gly-X-Y repeats, and a C-terminus. X or Y can be any amino acid except Gly, but are usually proline and hydroxyproline, respectively. b: The newly synthesized polypeptide chains are translocated into the lumen of the ER, where they undergo post-translational modifications. Hydroxylation of specific proline and lysine residues occurs, facilitated by enzymes such as prolyl hydroxylase and lysyl hydroxylase. c: In ECM, telopeptides in C and N terminals consist of amino acid sequences different from the rest of the collagen molecule are cleaved by proteases. d & e: Tropocollagens are packed end-to-end in a line and parallel aligned to form stable microfibril (94), which is cross-linked by covalent bonds that almost occurred between lysine and hydroxylysine mediated by lysyl oxidase.

## Appendix B: DMA test on a hard surface

Nano-DMA test on a hard, non-deformable glass slide as a baseline test. Due to glass’s high stiffness and negligible viscoelasticity, the test measures a very high storage modulus (representing the elastic response) with negligible loss modulus (representing viscous response), reflecting the rigid structure of glass. The phase angle between the excitation and response is zero degrees, indicating purely elastic behavior. This stable response provides a reference point for interpreting soft viscoelastic biological samples (here collagen) during comparative nano-rheometry testing.

**Figure B1:**
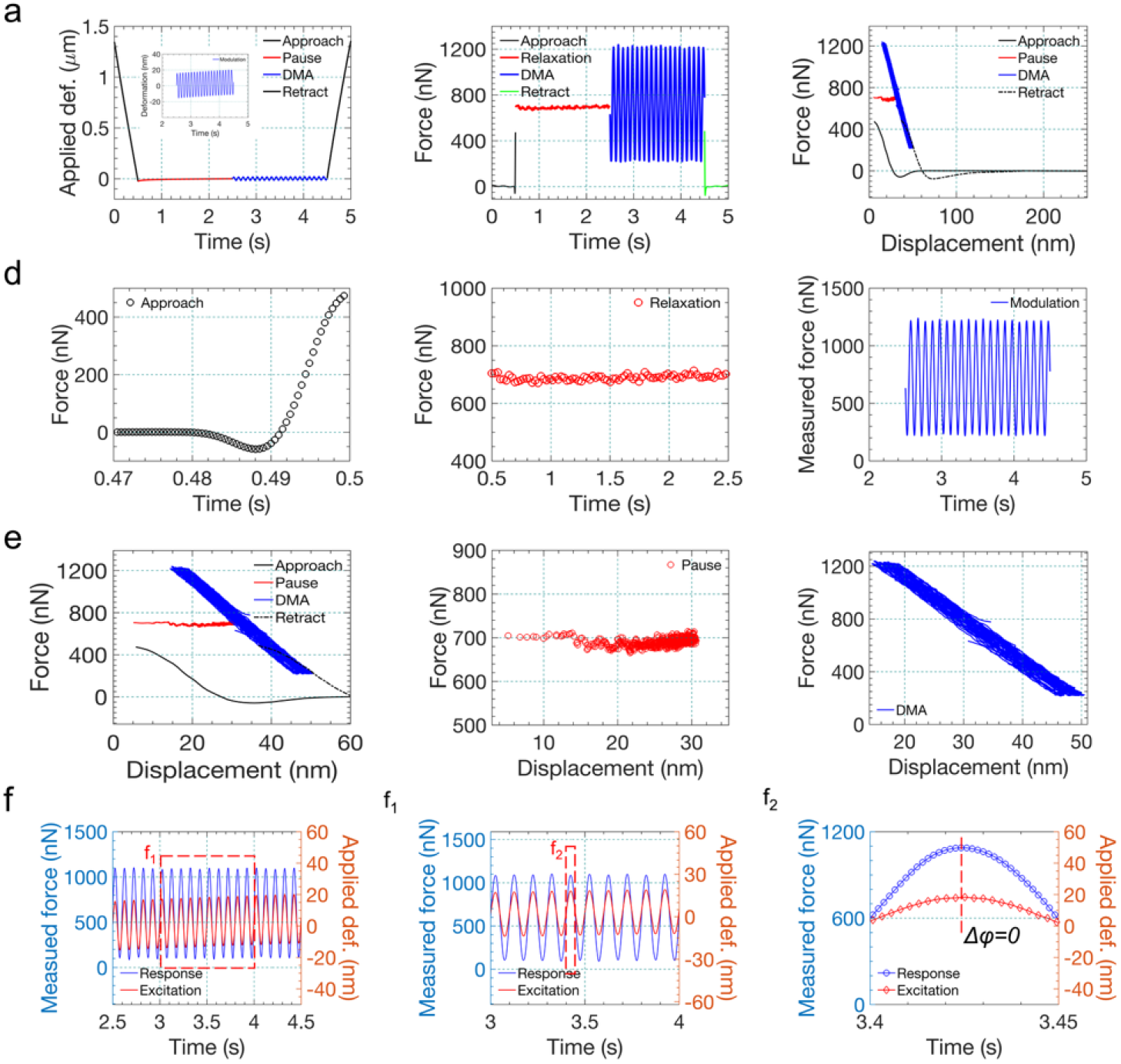
a: A typical Nano-DMA test was conducted on a glass slide, serving as a rigid surface, with varying loading phases depicted in panels b and c. Red graphs in panels d and e show no relaxation in time and Z-displacement fields respectively. As illustrated in panel f, there is zero phase difference between excitation and surface response for the glass surface.

## References

1. Buehler MJ. Nature designs tough collagen: explaining the nanostructure of collagen fibrils. Proceedings of the National Academy of Sciences. 2006;103(33):12285–90.

2. Fratzl P. Collagen: structure and mechanics, an introduction. Collagen: structure and mechanics: Springer; 2008. p. p1-13.

3. Manrique L, Moussa MS, Khan MT, Tahboub K, Ritchie RO, Asgari M, et al. Deformation of collagen-based tissues investigated using a systematic review and metaanalysis of synchrotron x-ray scattering studies. Cell Reports Physical Science. 2024.

4. Chattopadhyay S, Raines RT. Collagen-based biomaterials for wound healing. Biopolymers. 2014;101(8):821–33.

5. Yang W, Meyers MA, Ritchie RO. Structural architectures with toughening mechanisms in Nature: A review of the materials science of Type-I collagenous materials. Progress in Materials Science. 2019;103:425–83.

6. Vaez M, Asgari M, Hirvonen L, Bakir G, Khattignavong E, Ezzo M, et al. Modulation of the biophysical and biochemical properties of collagen by glycation for tissue engineering applications. Acta Biomaterialia. 2023;155:182–98.

7. Rufin M, Nalbach M, Rakuš M, Fuchs M, Poik M, Schitter G, et al. Methylglyoxal alters collagen fibril nanostiffness and surface potential. Acta Biomaterialia. 2024;189:208–16.

8. Asgari M, Latifi N, Giovanniello F, Espinosa HD, Amabili M. Revealing Layer-Specific Ultrastructure and Nanomechanics of Fibrillar Collagen in Human Aorta via Atomic Force Microscopy Testing: Implications on Tissue Mechanics at Macroscopic Scale. Advanced NanoBiomed Research. 2022;2(5):2100159.

9. Asgari M, Latifi N, Heris HK, Vali H, Mongeau L. In vitro fibrillogenesis of tropocollagen type III in collagen type I affects its relative fibrillar topology and mechanics. Scientific reports. 2017;7(1):1392.

10. Asgari M, Abi-Rafeh J, Hendy GN, Pasini D. Material anisotropy and elasticity of cortical and trabecular bone in the adult mouse femur via AFM indentation. Journal of the mechanical behavior of biomedical materials. 2019;93:81–92.

11. Amabili M, Asgari M, Breslavsky ID, Franchini G, Giovanniello F, Holzapfel GA. Microstructural and mechanical characterization of the layers of human descending thoracic aortas. Acta Biomaterialia. 2021;134:401–21.

12. Giovanniello F, Asgari M, Breslavsky ID, Franchini G, Holzapfel GA, Tabrizian M, et al. Development and mechanical characterization of decellularized scaffolds for an active aortic graft. Acta Biomaterialia. 2023;160:59–72.

13. Achterberg VF, Buscemi L, Diekmann H, Smith-Clerc J, Schwengler H, Meister J-J, et al. The nano-scale mechanical properties of the extracellular matrix regulate dermal fibroblast function. Journal of Investigative Dermatology. 2014;134(7):1862–72.

14. Viguet-Carrin S, Garnero P, Delmas P. The role of collagen in bone strength. Osteoporosis international. 2006;17:319–36.

15. Heino J. The collagen receptor integrins have distinct ligand recognition and signaling functions. Matrix Biology. 2000;19(4):319–23.

16. Latifi N, Asgari M, Vali H, Mongeau L. A tissue-mimetic nano-fibrillar hybrid injectable hydrogel for potential soft tissue engineering applications. Scientific reports. 2018;8(1):1047.

17. Guimarães CF, Gasperini L, Marques AP, Reis RL. The stiffness of living tissues and its implications for tissue engineering. Nature Reviews Materials. 2020;5(5):351–70.

18. Darling E, Zauscher S, Guilak F. Viscoelastic properties of zonal articular chondrocytes measured by atomic force microscopy. Osteoarthritis and cartilage. 2006;14(6):571–9.

19. Abi-Rafeh J, Asgari M, Troka I, Canaff L, Moussa A, Pasini D, et al. Genetic Deletion of Menin in Mouse Mesenchymal Stem Cells: An Experimental and Computational Analysis. JBMR plus. 2022;6(5):e10622.

20. Buehler MJ, Yung YC. Deformation and failure of protein materials in physiologically extreme conditions and disease. Nature materials. 2009;8(3):175–88.

21. Rigby BJ, Hirai N, Spikes JD, Eyring H. The mechanical properties of rat tail tendon. The Journal of general physiology. 1959;43(2):265–83.

22. Sasaki N, Shukunami N, Matsushima N, Izumi Y. Time-resolved X-ray diffraction from tendon collagen during creep using synchrotron radiation. Journal of biomechanics. 1999;32(3):285–92.

23. Wang XT, Ker RF. Creep rupture of wallaby tail tendons. Journal of experimental biology. 1995;198(3):831–45.

24. Ghodsi H, Darvish K. Investigation of mechanisms of viscoelastic behavior of collagen molecule. Journal of the mechanical behavior of biomedical materials. 2015;51:194–204.

25. Ghodsi H, Darvish K. Characterization of the viscoelastic behavior of a simplified collagen micro-fibril based on molecular dynamics simulations. Journal of the mechanical behavior of biomedical materials. 2016;63:26–34.

26. Gautieri A, Vesentini S, Redaelli A, Buehler MJ. Viscoelastic properties of model segments of collagen molecules. Matrix Biology. 2012;31(2):141–9.

27. Andriotis OG, Nalbach M, Thurner PJ. Mechanics of isolated individual collagen fibrils. Acta Biomaterialia. 2023;163:35–49.

28. Liu Y, Ballarini R, Eppell SJ. Tension tests on mammalian collagen fibrils. Interface focus. 2016;6(1):20150080.

29. Shen ZL, Dodge MR, Kahn H, Ballarini R, Eppell SJ. In vitro fracture testing of submicron diameter collagen fibril specimens. Biophysical journal. 2010;99(6):1986–95.

30. Shen ZL, Dodge MR, Kahn H, Ballarini R, Eppell SJ. Stress-strain experiments on individual collagen fibrils. Biophysical journal. 2008;95(8):3956–63.

31. Shen ZL, Kahn H, Ballarini R, Eppell SJ. Viscoelastic properties of isolated collagen fibrils. Biophysical journal. 2011;100(12):3008–15.

32. Tang Y, Ballarini R, Buehler MJ, Eppell SJ. Deformation micromechanisms of collagen fibrils under uniaxial tension. Journal of The Royal Society Interface. 2010;7(46):839–50.

33. Yang F, Das D, Karunakaran K, Genin GM, Thomopoulos S, Chasiotis I. Nonlinear time-dependent mechanical behavior of mammalian collagen fibrils. Acta Biomaterialia. 2023;163:63–77.

34. Gautieri A, Vesentini S, Redaelli A, Ballarini R. Modeling and measuring visco-elastic properties: From collagen molecules to collagen fibrils. International Journal of Non-Linear Mechanics. 2013;56:25–33.

35. Sasaki N, Odajima S. Elongation mechanism of collagen fibrils and force-strain relations of tendon at each level of structural hierarchy. Journal of biomechanics. 1996;29(9):1131–6.

36. Sasaki N, Odajima S. Stress-strain curve and Young’s modulus of a collagen molecule as determined by the X-ray diffraction technique. Journal of biomechanics. 1996;29(5):655–8.

37. Gupta H, Seto J, Krauss S, Boesecke P, Screen H. In situ multi-level analysis of viscoelastic deformation mechanisms in tendon collagen. Journal of structural biology. 2010;169(2):183–91.

38. Sasaki N, Nakayama Y, Yoshikawa M, Enyo A. Stress relaxation function of bone and bone collagen. Journal of biomechanics. 1993;26(12):1369–76.

39. Yang L, Van der Werf K, Dijkstra PJ, Feijen J, Bennink ML. Micromechanical analysis of native and cross-linked collagen type I fibrils supports the existence of microfibrils. Journal of the mechanical behavior of biomedical materials. 2012;6:148–58.

40. Van Der Rijt JA, Van Der Werf KO, Bennink ML, Dijkstra PJ, Feijen J. Micromechanical testing of individual collagen fibrils. Macromolecular bioscience. 2006;6(9):697–702.

41. Svensson RB, Hassenkam T, Hansen P, Magnusson SP. Viscoelastic behavior of discrete human collagen fibrils. Journal of the Mechanical Behavior of Biomedical Materials. 2010;3(1):112–5.

42. Svensson RB, Mulder H, Kovanen V, Magnusson SP. Fracture mechanics of collagen fibrils: influence of natural cross-links. Biophysical journal. 2013;104(11):2476–84.

43. Yang L, van der Werf KO, Koopman BF, Subramaniam V, Bennink ML, Dijkstra PJ, et al. Micromechanical bending of single collagen fibrils using atomic force microscopy. Journal of Biomedical Materials Research Part A. 2007;82(1):160–8.

44. Yang L, Van der Werf KO, Fitié CF, Bennink ML, Dijkstra PJ, Feijen J. Mechanical properties of native and cross-linked type I collagen fibrils. Biophysical journal. 2008;94(6):2204–11.

45. Grant CA, Brockwell DJ, Radford SE, Thomson NH. Tuning the elastic modulus of hydrated collagen fibrils. Biophysical journal. 2009;97(11):2985–92.

46. Grant CA, Brockwell DJ, Radford SE, Thomson NH. Effects of hydration on the mechanical response of individual collagen fibrils. Applied Physics Letters. 2008;92(23).

47. Heim AJ, Matthews WG, Koob TJ. Determination of the elastic modulus of native collagen fibrils via radial indentation. Applied physics letters. 2006;89(18).

48. Grant CA, Phillips MA, Thomson NH. Dynamic mechanical analysis of collagen fibrils at the nanoscale. Journal of the mechanical behavior of biomedical materials. 2012;5(1):165–70.

49. Andriotis OG, Manuyakorn W, Zekonyte J, Katsamenis OL, Fabri S, Howarth PH, et al. Nanomechanical assessment of human and murine collagen fibrils via atomic force microscopy cantilever-based nanoindentation. Journal of the mechanical behavior of biomedical materials. 2014;39:9–26.

50. Minary-Jolandan M, Yu M-F. Nanomechanical heterogeneity in the gap and overlap regions of type I collagen fibrils with implications for bone heterogeneity. Biomacromolecules. 2009;10(9):2565–70.

51. Baldwin SJ, Kreplak L, Lee JM. Characterization via atomic force microscopy of discrete plasticity in collagen fibrils from mechanically overloaded tendons: Nano-scale structural changes mimic rope failure. Journal of the mechanical behavior of biomedical materials. 2016;60:356–66.

52. Baldwin SJ, Quigley AS, Clegg C, Kreplak L. Nanomechanical mapping of hydrated rat tail tendon collagen I fibrils. Biophysical journal. 2014;107(8):1794–801.

53. Baldwin SJ, Sampson J, Peacock CJ, Martin ML, Veres SP, Lee JM, et al. A new longitudinal variation in the structure of collagen fibrils and its relationship to locations of mechanical damage susceptibility. Journal of the mechanical behavior of biomedical materials. 2020;110:103849.

54. Gisbert VG, Benaglia S, Uhlig MR, Proksch R, Garcia R. High-speed nanomechanical mapping of the early stages of collagen growth by bimodal force microscopy. ACS nano. 2021;15(1):1850–7.

55. Egan JM. A constitutive model for the mechanical behaviour of soft connective tissues. Journal of biomechanics. 1987;20(7):681–92.

56. Haut RC, Little RW. A constitutive equation for collagen fibers. Journal of biomechanics. 1972;5(5):423–30.

57. Peña E, Pena J, Doblaré M. On modelling nonlinear viscoelastic effects in ligaments. Journal of biomechanics. 2008;41(12):2659–66.

58. Puxkandl R, Zizak I, Paris O, Keckes J, Tesch W, Bernstorff S, et al. Viscoelastic properties of collagen: synchrotron radiation investigations and structural model. Philosophical Transactions of the Royal Society of London Series B: Biological Sciences. 2002;357(1418):191–7.

59. Woo S-Y, Johnson GA, Smith BA. Mathematical modeling of ligaments and tendons. 1993.

60. Fung Y-C, Skalak R. Biomechanics. Mechanical properties of living tissues. Journal of Applied Mechanics. 1982;49(2):464.

61. Silver FH, Ebrahimi A, Snowhill PB. Viscoelastic properties of self-assembled type I collagen fibers: molecular basis of elastic and viscous behaviors. Connective Tissue Research. 2002;43(4):569–80.

62. Sopakayang R, De Vita R, Kwansa A, Freeman JW. Elastic and viscoelastic properties of a type I collagen fiber. Journal of Theoretical Biology. 2012;293:197–205.

63. Efremov YM, Wang W-H, Hardy SD, Geahlen RL, Raman A. Measuring nanoscale viscoelastic parameters of cells directly from AFM force-displacement curves. Scientific reports. 2017;7(1):1541.

64. Kolluru PV, Eaton MD, Collinson DW, Cheng X, Delgado DE, Shull KR, et al. AFM-based dynamic scanning indentation (DSI) method for fast, high-resolution spatial mapping of local viscoelastic properties in soft materials. Macromolecules. 2018;51(21):8964–78.

65. Collinson DW, Sheridan RJ, Palmeri MJ, Brinson LC. Best practices and recommendations for accurate nanomechanical characterization of heterogeneous polymer systems with atomic force microscopy. Progress in Polymer Science. 2021;119:101420.

66. Abuhattum S, Kotzbeck P, Schlüßler R, Harger A, Ariza de Schellenberger A, Kim K, et al. Adipose cells and tissues soften with lipid accumulation while in diabetes adipose tissue stiffens. Scientific Reports. 2022;12(1):10325.

67. Efremov YM, Shpichka A, Kotova S, Timashev P. Viscoelastic mapping of cells based on fast force volume and PeakForce Tapping. Soft Matter. 2019;15(27):5455–63.

68. Ting T. The contact stresses between a rigid indenter and a viscoelastic half-space. 1966.

69. Garcia PD, Garcia R. Determination of the viscoelastic properties of a single cell cultured on a rigid support by force microscopy. Nanoscale. 2018;10(42):19799–809.

70. Alcaraz J, Buscemi L, Grabulosa M, Trepat X, Fabry B, Farré R, et al. Microrheology of human lung epithelial cells measured by atomic force microscopy. Biophysical journal. 2003;84(3):2071–9.

71. Kollmannsberger P, Fabry B. Active soft glassy rheology of adherent cells. Soft Matter. 2009;5(9):1771–4.

72. Efremov YM, Okajima T, Raman A. Measuring viscoelasticity of soft biological samples using atomic force microscopy. Soft matter. 2020;16(1):64–81.

73. Hiratsuka S, Mizutani Y, Tsuchiya M, Kawahara K, Tokumoto H, Okajima T. The number distribution of complex shear modulus of single cells measured by atomic force microscopy. Ultramicroscopy. 2009;109(8):937–41.

74. Rother J, Nöding H, Mey I, Janshoff A. Atomic force microscopy-based microrheology reveals significant differences in the viscoelastic response between malign and benign cell lines. Open biology. 2014;4(5):140046.

75. Efremov YM, Dokrunova A, Bagrov D, Kudryashova K, Sokolova O, Shaitan K. The effects of confluency on cell mechanical properties. Journal of biomechanics. 2013;46(6):1081–7.

76. Dufrêne YF, Pelling AE. Force nanoscopy of cell mechanics and cell adhesion. Nanoscale. 2013;5(10):4094–104.

77. Nair AK, Gautieri A, Chang S-W, Buehler MJ. Molecular mechanics of mineralized collagen fibrils in bone. Nature communications. 2013;4(1):1724.

78. Weiner S, Wagner HD. The material bone: structure-mechanical function relations. Annual review of materials science. 1998;28(1):271–98.

79. Robinson R, Watson M. Collagen-crystal relationships in bone as seen in the electron microscope. The anatomical record. 1952;114(3):383–409.

80. Orgel JP, Irving TC, Miller A, Wess TJ. Microfibrillar structure of type I collagen in situ. Proceedings of the National Academy of Sciences. 2006;103(24):9001–5.

81. Tang M, Li T, Gandhi NS, Burrage K, Gu Y. Heterogeneous nanomechanical properties of type I collagen in longitudinal direction. Biomechanics and modeling in mechanobiology. 2017;16:1023–33.

82. Herchenhan A, Uhlenbrock F, Eliasson P, Weis M, Eyre D, Kadler KE, et al. Lysyl oxidase activity is required for ordered collagen fibrillogenesis by tendon cells. Journal of Biological Chemistry. 2015;290(26):16440–50.

83. Kong W, Lyu C, Liao H, Du Y. Collagen crosslinking: effect on structure, mechanics and fibrosis progression. Biomedical Materials. 2021;16(6):062005.

84. Wilson SL, Guilbert M, Sulé-Suso J, Torbet J, Jeannesson P, Sockalingum GD, et al. A microscopic and macroscopic study of aging collagen on its molecular structure, mechanical properties, and cellular response. The FASEB Journal. 2014;28(1):14–25.

85. Mohammadkhah M, Klinge S. The importance of consideration of collagen crosslinks in computational models of collagen-based tissues. Journal of the Mechanical Behavior of Biomedical Materials. 2023;148:106203.

86. Shoulders MD, Raines RT. Collagen structure and stability. Annual review of biochemistry. 2009;78:929–58.

87. Parry DA. The molecular fibrillar structure of collagen and its relationship to the mechanical properties of connective tissue. Biophysical chemistry. 1988;29(1-2):195–209.

88. Ottani V, Martini D, Franchi M, Ruggeri A, Raspanti M. Hierarchical structures in fibrillar collagens. Micron. 2002;33(7-8):587–96.

89. Vassaux M. Heterogeneous structure and dynamics of water in a hydrated collagen microfibril. Biomacromolecules. 2024.

90. Ghanaeian A, Soheilifard R. Mechanical elasticity of proline-rich and hydroxyproline-rich collagen-like triple-helices studied using steered molecular dynamics. Journal of the Mechanical Behavior of Biomedical Materials. 2018;86:105–12.

91. Naomi R, Ridzuan PM, Bahari H. Current insights into collagen type I. Polymers. 2021;13(16):2642.

92. Ling S, Chen W, Fan Y, Zheng K, Jin K, Yu H, et al. Biopolymer nanofibrils: Structure, modeling, preparation, and applications. Progress in polymer science. 2018;85:1–56.

93. Brazel D, Oberbäumer I, Dieringer H, Babel W, Glanville RW, Deutzmann R, et al. Completion of the amino acid sequence of the α1 chain of human basement membrane collagen (type IV) reveals 21 non-triplet interruptions located within the collagenous domain. European journal of biochemistry. 1987;168(3):529–36.

94. Abou Neel EA, Bozec L, Knowles JC, Syed O, Mudera V, Day R, et al. Collagen— emerging collagen based therapies hit the patient. Advanced drug delivery reviews. 2013;65(4):429–56.

95. Ricard-Blum S. The collagen family. Cold Spring Harbor perspectives in biology. 2011;3(1):a004978.

96. Myllyharju J. Intracellular post-translational modifications of collagens. Collagen: primer in structure, processing and assembly. 2005:115–47.

97. Myllyharju J, Kivirikko KI. Collagens, modifying enzymes and their mutations in humans, flies and worms. TRENDS in Genetics. 2004;20(1):33–43.

98. Stefanovic B. RNA protein interactions governing expression of the most abundant protein in human body, type I collagen. Wiley Interdisciplinary Reviews: RNA. 2013;4(5):535–45.

99. Xu S, Xu H, Wang W, Li S, Li H, Li T, et al. The role of collagen in cancer: from bench to bedside. Journal of translational medicine. 2019;17:1–22.

100. Zhao C, Xiao Y, Ling S, Pei Y, Ren J. Structure of collagen. Fibrous Proteins: Design, Synthesis, and Assembly. 2021:17–25.

101. Lucero H, Kagan H. Lysyl oxidase: an oxidative enzyme and effector of cell function. Cellular and Molecular Life Sciences CMLS. 2006;63:2304–16.

102. Molnar J, Fong K, He Q, Hayashi K, Kim Y, Fong S, et al. Structural and functional diversity of lysyl oxidase and the LOX-like proteins. Biochimica et biophysica acta (BBA)-proteins and proteomics. 2003;1647(1-2):220–4.

103. Kang AH, Gross J. Relationship between the Intra and Inter molecular Cross-links of Collagen. Proceedings of the National Academy of Sciences. 1970;67(3):1307–14.

104. Fratzl P, Weinkamer R. Nature’s hierarchical materials. Progress in materials Science. 2007;52(8):1263–334.

