## Supplementary information for "Ultrastructural viscoelastic behavior of fibrillar collagen identified by AFM Nano-Rheometry and direct indentation"

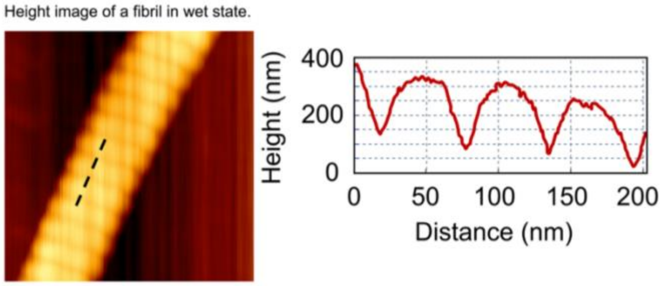


Supplementary Figure 1: Height topography of an isolated fibril in wet state. While the gap/overlap regions are distinguishable, the D-banding is less apparent in the hydrated state than in dry state.

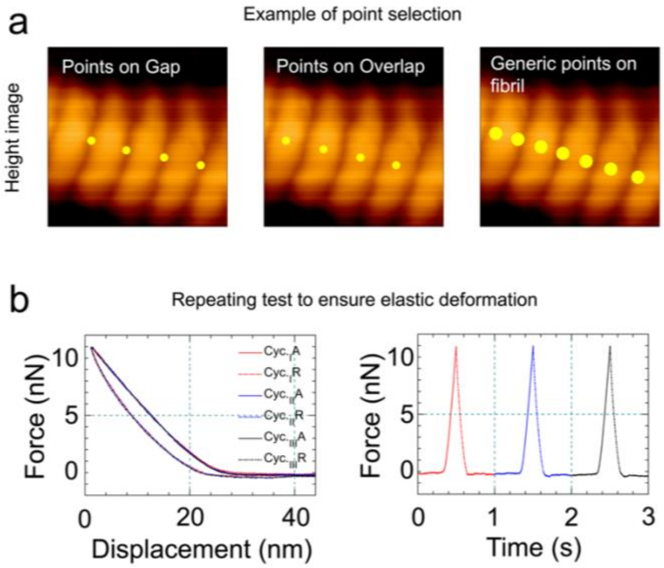


Supplementary Figure 2: (a) Representative example illustrating the point selection strategy used to differentiate between the gap and overlap subregions of a collagen fibril compared to the entire fibril. Although the D-banding appears less pronounced in the hydrated state, it is essential to select indentation points under fluid conditions. Indentation measurements of the subregions were performed using an AFM tip with a smaller radius (R=20 nm) to achieve higher spatial resolution and minimize lateral interference between adjacent domains. In contrast, indentation of the whole fibril was conducted using a larger AFM tip (R=50 nm), allowing for averaged mechanical property assessment over a broader contact area. (b) Verification of elastic deformation through repeated indentation at the same location. The indentation procedure was repeated three times under identical loading conditions to ensure that the mechanical response remained within the elastic regime, and no plastic or permanent deformation occurred. The overlapping force-displacement curves from the three cycles confirm purely elastic behavior, as indicated by the complete recovery of the fibril after each indentation cycle.

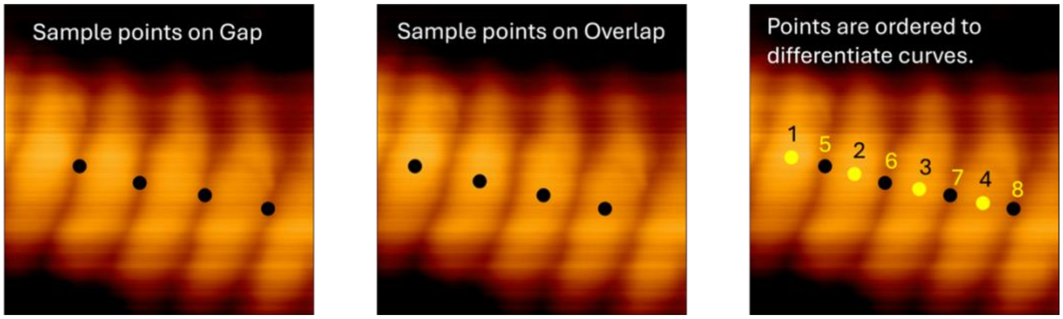


**Supplementary Figure 2: Strategy for Recording Indentation Data.** The indentation procedure begins with the systematic selection of measurement points on one subregion of the collagen fibril, typically the overlap region, followed by the selection of points on the second subregion, such as the gap region. Each indentation point is clearly indexed, taking into account both the total number of locations and the number of repeated measurements at each point. This indexing allows for precise tracking and identification of individual force-displacement curves. All indentation data are recorded sequentially and saved in a structured format. Post-acquisition, the data are organized and categorized according to the specific subregion (gap or overlap), enabling accurate comparative mechanical analysis between the two subregions.
